# Poor matching of action codes challenges ‘mirroring’ in macaque F5 mirror neurons

**DOI:** 10.1101/2022.02.01.478703

**Authors:** Joern K. Pomper, Mohammad Shams-Ahmar, Shengjun Wen, Friedemann Bunjes, Peter Thier

## Abstract

The discharge of F5 mirror neurons of a monkey observing another individual performing an action has been interpreted as a motor representation, stimulated by sensory input, ultimately serving action understanding. This hypothesis requires mirror neurons to exhibit an action tuning that is matched between action observation and execution. However, the evidence for this is insufficient, mainly because previous work has suffered from a limited exploration of action space tuning and an object vs. action vision confound. To overcome these limitations, we conducted an experiment in which identical objects had to be manipulated in three different ways in order to serve distinct action goals. We show that the population of F5 mirror neurons can dissociate the three actions almost perfectly, during both observation and execution. However, the population code for action execution was only poorly matched to the code for observation. Just a few neurons exhibited only matched codes, yet, always confined to short, varying segments of the overall action. These findings challenge the hypothesis that an observer understands the actions of others by the activation of motor representations of the observed actions. Rather they support the alternative notion that F5 might use information on observed actions to select behavioral responses.

## Introduction

Mirror neurons have been defined as neurons that modulate their discharge rate not only in response to the execution of an action directed toward an object but also in response to the observation of a similar action carried out by others (Gallese et al., 1996; Rizzolatti et al., 1996). The discovery of neurons characterized by this intriguing combination of response preferences in premotor cortical area F5 of macaque monkeys (di Pellegrino et al., 1992) attracted great interest, which resulted not only in a large number of subsequent studies in which the properties of these neurons were explored in more detail (Rizzolatti et al., 2014) but also in the discovery of mirror neurons in other cortical areas of monkeys (Fogassi et al., 2005; Papadourakis and Raos, 2019) and species, including humans (Mukamel et al., 2010).

Down to the present day, researchers studying mirror neurons have been mesmerized by the possibility that these peculiar neurons might play a role in processing information on the other’s actions, rather than being confined to action planning and/or control, the latter perhaps suggested by their occurrence in area F5, formerly thought to be confined to motor planning (Rizzolatti et al., 1988). Nevertheless, why should information on the other’s action be processed by neurons in a motor area? A possible answer is that the observation of others’ action might resonate in a neuronal machinery, at other times controlling similar actions of the observer. In other words, action understanding is thought to arise from allowing the observer’s action planning and execution machinery to simulate the other’s action (Rizzolatti et al., 1996).

Simulation of others’ action based on F5 mirror neurons would require that the activity of these neurons is the same (or similar enough), no matter if ignited by the desire to engage in a particular action or in its observation. This is the tenet of the *mirror mechanism* (Rizzolatti and Sinigaglia, 2016). Although some studies suggested that indeed a substantial fraction of mirror neurons might satisfy this hypothesis (di Pellegrino et al., 1992; Gallese et al., 1996; Rizzolatti et al., 1996), others have remained skeptical, considering inconsistencies and limitations of the experimental evidence available that may not suffice to support the strong requirements of the *mirror mechanism* (Cook and Bird, 2013; Csibra, 2007; Hickok, 2009; Thompson et al., 2019). For instance, whereas Gallese and colleagues found that about a third of mirror neurons exhibited the same grip type preferences during execution and observation (Gallese et al., 1996), a recent study reported similar grip type tuning during execution and observation tasks only at the population level. However, in the latter study, the relationship of grip type preferences for execution and observation at the single-cell level was reported to be random (Papadourakis and Raos, 2019). The lack of consistent evidence for mirroring grip type preferences might be because mirror neurons serving the assumed *mirror mechanism* may not care about specific grip types but about action goals, independent of a consideration of low-level features that matter in the description of specific grip types (Rizzolatti and Sinigaglia, 2010). In accordance with this possibility, previous studies could indeed show that the response of some mirror neurons to executed or observed actions can be largely independent of the effector used — hand, mouth, or tools including reverse pliers — as long as the action goal was maintained (Ferrari et al., 2005; Rizzolatti and Sinigaglia, 2016; Rochat et al., 2010).

Encouraged by these results, it was concluded that mirror neurons may indeed encode an action goal as basis of action understanding (Rizzolatti and Sinigaglia, 2016). However, while certainly intriguing, this conclusion suffers from a shortage of expedient experimental data needed to critically assess it. For instance, responses to execution and to observation should not only match for the preferred goal identified by the strongest modulation but also for others, albeit giving rise to less discharge modulation. In order to test this prediction we decided to carry out a rigorous comparison of responses to three different action goals by asking the experimental animal to manipulate a visually identical object in three different ways, namely to lift it, to twist it, or to shift it. This enabled us to characterize an action tuning, while avoiding a confounding influence of different objects. The actions differed in the ultimate action goal given by different end positions and the type of manipulation requiring different grip types. We reasoned that if mirror neurons were the substrate of a *mirror mechanism* supporting a representation of action goals, then this experimental paradigm should reveal it. And since the *mirror mechanism* demands a matched code between action execution and observation, it should be possible to decode the specific action from the activity of mirror neurons during observation if we knew how the same action was encoded during execution.

We found that population-based decoders, trained on action observation trials predicted observed actions with a reliability of up to 100% in dissociating the three different action types. However, decoders trained on execution trials predicted observed actions only poorly (up to 55%; chance level 33.3%). Even in a small subpopulation, in which execution-trained decoders predicted observed actions equally well as observation-trained decoders, this prediction was confined to a segment of the overall action epoch. Hence, these results prompt a reconsideration of the role of mirror neurons beyond the tight bounds of a *mirror mechanism*.

## Results

Two rhesus macaques were trained to perform the behavioral paradigm depicted in figure 1. The paradigm consisted of two tasks that were performed in two separate blocks (Fig. 1A): an execution task, in which the experimental monkey had to perform one out of three possible actions on objects with identical visual appearance (Fig. 1B, top row), and an observation task, in which he was asked to pay attention to the same actions carried out by a human actor in front of him (Fig. 1B, bottom row). Which action the object at stake afforded was indicated by a color cue provided by an LED next to the respective target object (Fig. 1A). The target object was always the object coming to a hold right in front of the respective actor (the monkey or the human) after an initial random rotation (in darkness) of the table carrying the three identically looking objects, carried out in order to ensure that memories of previous action type choices were useless. Distinct events allowed the separation of six epochs (Fig. 1C).

**Figure 1.**
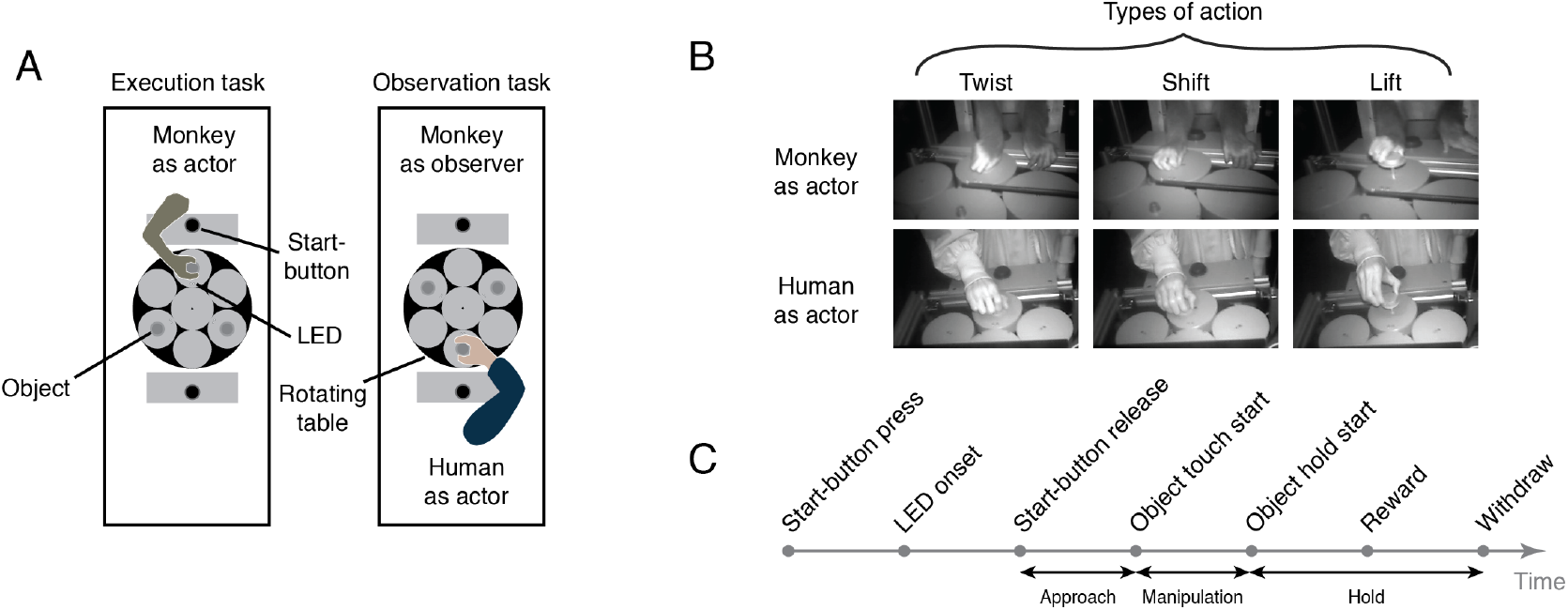
The behavioral paradigm. A) Experimental setup with three identical objects positioned on a rotating table in front of the actor. Each object could be acted on in only one way. The type of action required was cued by the color of an LED next to the object. After LED onset, the actor was allowed to release the start-button. B) Photographs of the three actions in execution and observation tasks at the time when the object was held in its target position. C) The sequence of events in a trial.

Out of 240 neurons recorded from two monkeys, 177 (74%) met the mirror neuron criteria outlined in the Methods and were used for the following analyses. As summarized in Table 1, the time between two adjacent events varied depending on the trial, type of action, and task. In order to make the timing of events comparable, we transformed the absolute times between events to a relative time by dividing the time interval between every two adjacent events into quartiles. Hence, each trial was divided into 24 time bins between the time of pressing the start-button and the time of withdrawing from the object.

**Table 1.**
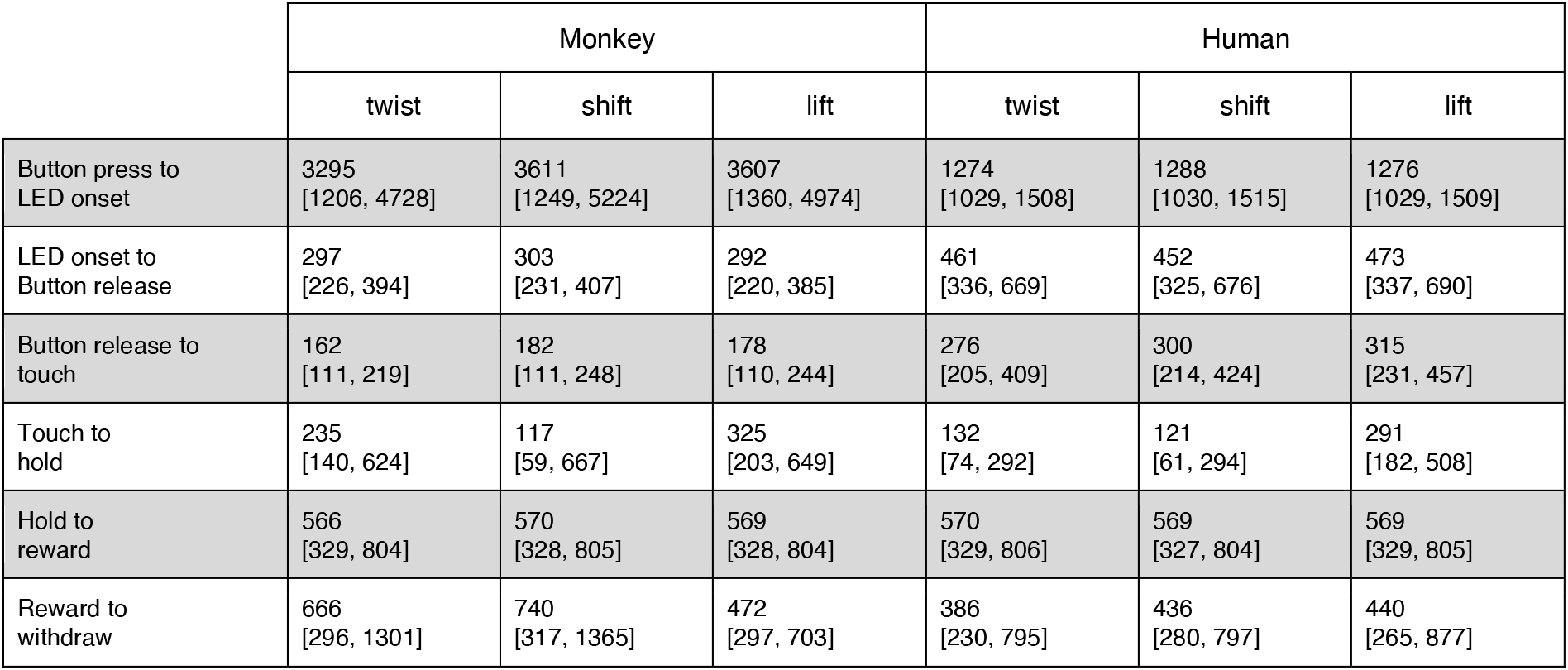
The average duration (in ms) across trials of each task period per action for monkeys and humans. The lower and upper bounds of the 95% confidence intervals are shown in brackets.

### Only a minority of mirror neurons exhibits congruent action tuning in at least one time bin

Similar to previous approaches to mirror neurons, we identified the preferred action of a neuron as the single action or a pair of actions (in case the two actions were not statistically different) that yielded a discharge rate that was significantly larger than those associated with other action type(s) (see Methods). We assessed the preferences independently for each of the 24 bins. In the execution task, most neurons had a preferred action in at least one time bin (n = 137, 77%). In the observation task, this proportion was slightly less (n = 104, 59%). In total, it was common that the preferred action differed across the time bins (see the exemplary neurons in fig. 2).

**Figure 2.**
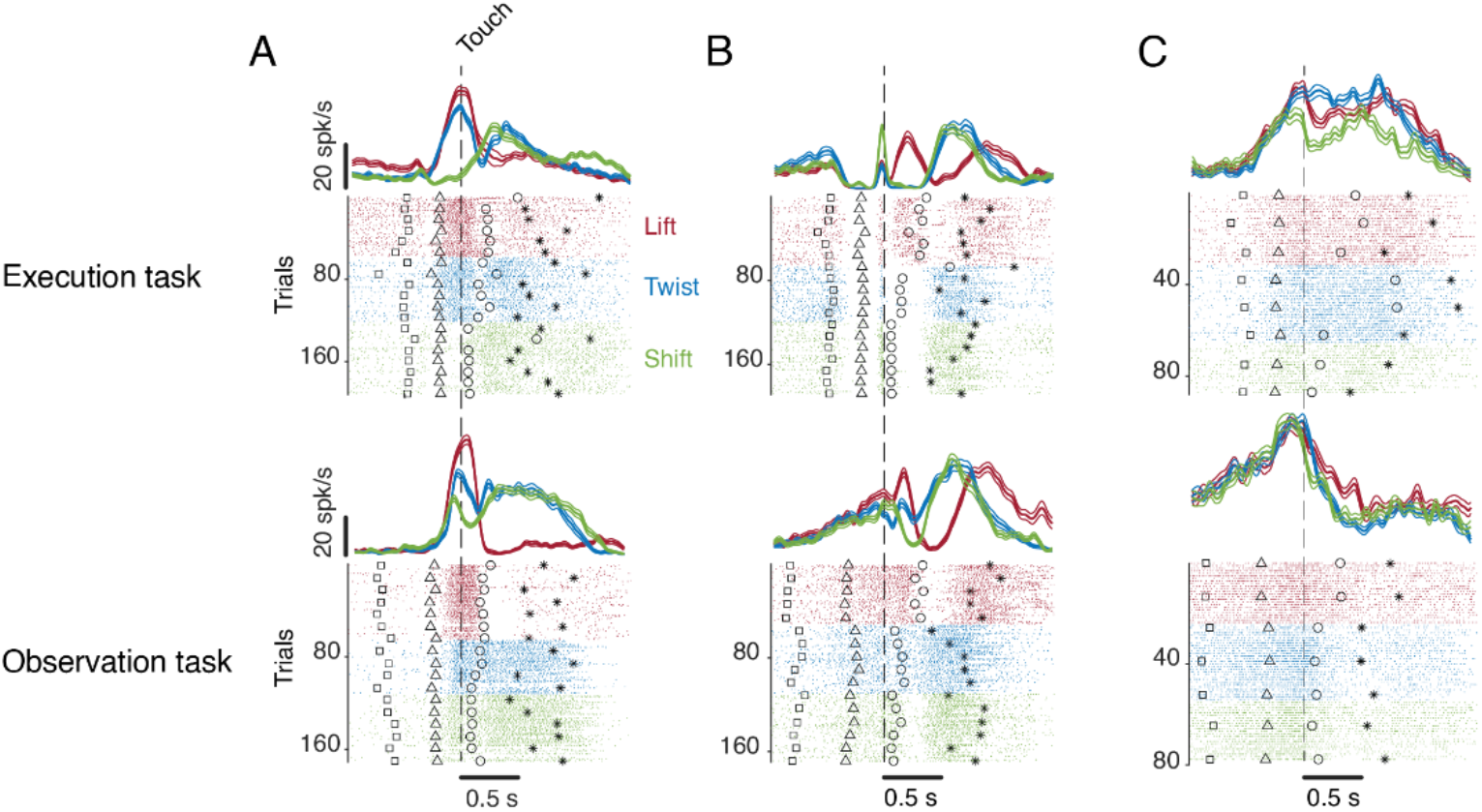
Exemplary F5 mirror neurons. Spike density functions and raster plots are aligned to the time of touch start (vertical dashed line). A) A neuron that prefers a lift at the time of touch and then later a twist and a shift in both tasks. B) A neuron that prefers a shift during execution and a lift during observation at the time of touch, but a lift after touch in both tasks. C) A neuron that has no or different preference(s) during execution and observation. In A-C, markers indicate the time of four events around the time of touch: LED onset (square), start-button release (triangle), object hold start (circle), and reward (star). For better visualization, the events are shown for only about 10% of trials.

The comparison of action preferences between execution and observation showed that a quarter of all mirror neurons (n = 41, 23%) had the same action preference in at least one time bin. Some neurons maintained the same preference across multiple time bins (such as the neurons depicted in Fig. 2A), whereas others had the same preference in only one time bin (such as the one in Fig. 2B). The remaining neurons that had a preference in at least one time bin during observation (n = 63, 36%) had either no preference or a different preference in that time bin during execution (such as Fig. 2C). In short, only a minority of mirror neurons exhibited the same action preferences for execution and observation. In these neurons, however, congruent preferences were usually confined to restricted periods of time.

### The action encoding in the execution and observation tasks transfers only incompletely to the respective other task

The question is if an ideal observer, having to consider all mirror neurons rather than being able to select the minority exhibiting congruent action preferences (as indicated by the preliminary analysis), would still be able to predict the type of action carried out? In order to address this question, we resorted to a linear discriminant classification of actions, allowing us to determine if the neural code of actions in the execution task would be able to predict the actions during observation. In a first step, we asked how well the three actions could be decoded during the execution task. We trained a classifier for each neuron and time bin based on 75% of the execution trials (as described in more detail in the Methods) and examined how well it predicted the action in the remaining trials (exe → exe, blue curve in Fig. 3A). We found that the mean performance of the classifiers calculated across all mirror neurons increased sharply in the third time bin after LED onset (Fig. 3A). This increase continued with a lower slope until it reached its maximum of about 42% successful discrimination (chance level 33.3%) in the first time bin after the object had been moved to its final position.

**Figure 3.**
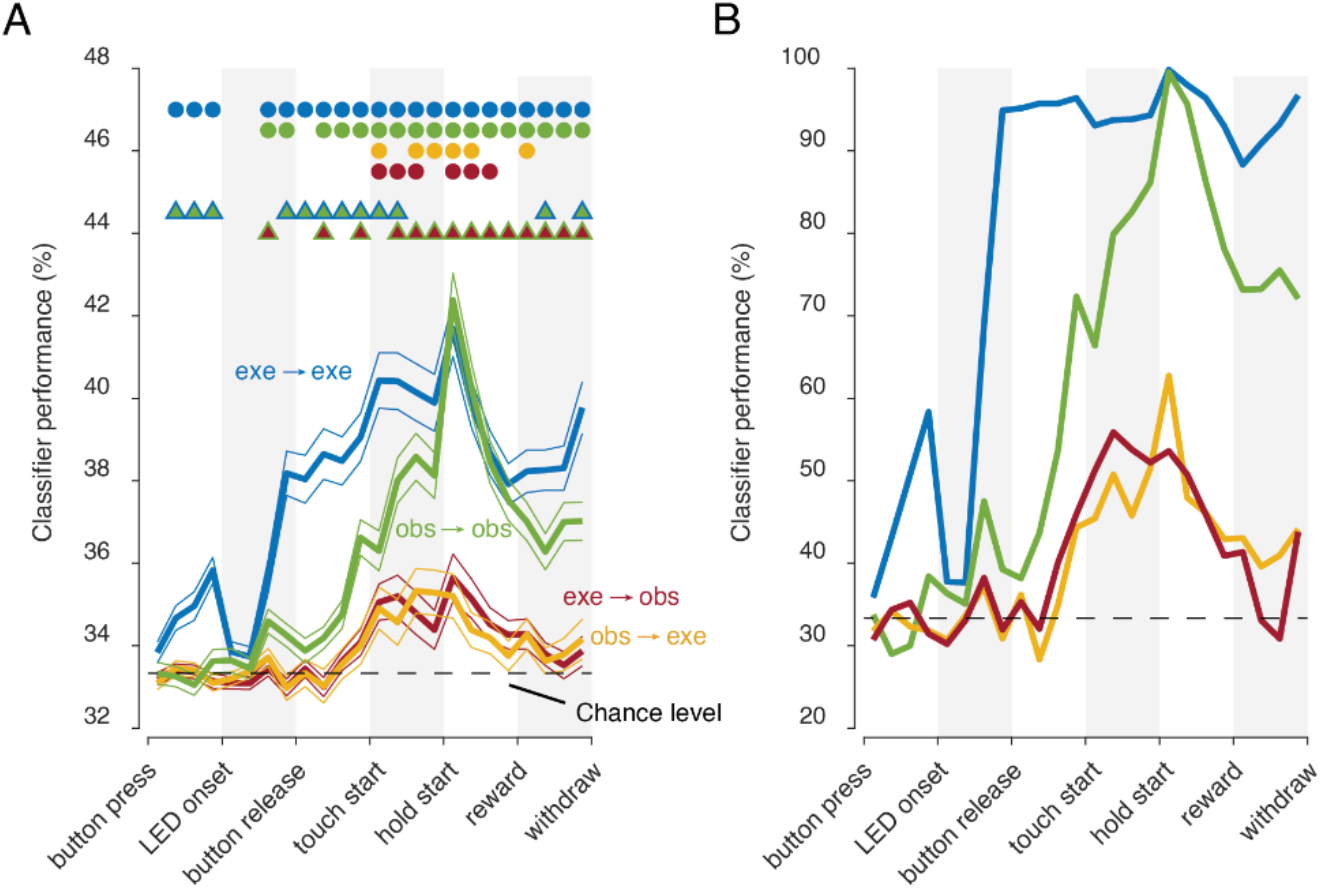
Classification of the three actions. A) Single neuron classification results (mean ± s.e.m, n = 177 neurons) per time bin, execution-trained classifiers tested with execution trials (exe → exe, blue), tested with observation trials (exe → obs, red). Observation-trained classifiers tested with observation trials (obs → obs, green), and execution trials (obs → exe, orange). Colored dots indicate time bins with significant performance above chance (Benjamini-Hochberg-corrected one-sided signed-rank tests). Two colored triangles indicate time bins with significant differences in the performance between the corresponding colors (Benjamini-Hochberg-corrected two-sided signed-rank tests). Shaded gray areas are for better visual dissociation of the task periods, each consisting of four time bins. B) Population classification results per time bin. Same as A, but here, all neurons (n = 177) constructed the 177 features of a classifier.

In a second step, we asked how well the three actions could be decoded during the observation task. We trained a classifier for each neuron and time bin based on 75% of the observation trials and examined how well it predicted the action from the remaining trials (obs → obs, green curve in Fig. 3A). The rise of discrimination performance started three time bins later, in the second time bin after start-button release. However, the level of maximal discrimination achieved (~42%) was comparable to the execution decoder’s performance and the peak was reached at the same time. In sum, the actions could be read out from the discharge rate of mirror neurons during both execution and observation tasks equally well at least in one time bin after the object had been moved to its final position.

We finally asked if the encoding of the actions during the execution and observation tasks matched. Therefore, we applied the classifiers trained in one task to the other task respectively (exe → obs, the red curve in Fig. 3A; obs → exe, the orange curve in Fig. 3A). Regardless of the direction of the test (exe → obs or obs → exe), we observed an increase in the discrimination performance that started in the third time bin after the start-button release and reached the highest level of performance while the manipulation was carried out. However, the maximum level of performance attained was only ~36% for both transfer directions, which was significantly below the exe → exe and the obs → obs discrimination performances. Hence, there is a match between the codes for the three actions in the execution and observation tasks, yet the match is poor.

A biologically plausible way to read out actions offered by a population of mirror neurons would be to rely on the collective activity of all, rather than on individual neurons. Therefore we trained a linear discrimination classifier on the discharge rates of all 177 mirror neurons, separately for execution and observation, in a bin-wise manner, fully analogous to the procedure deployed for the single neuron classification. As shown in figure 3B, we found a very high performance for both the exe → exe and the obs → obs classifications with a time course comparable to the single neuron-based results. Shortly after reaching the final position of the object, the performance was optimal (100%) for both the exe → exe and the obs → obs classifications. The classifiers trained on data from one task and tested on data from the other task gave a much weaker performance of about 55% to 60%. In sum, as expected, the population classifiers performed much better than the single neuron classifiers. However, similar to the single neuron classifications, the across-task performances were consistently below the within-task performances.

### There is a cluster of time bins with a matched code

It is conceivable that the across-task performances of all single neurons were consistently (across neurons and time bins) weaker than the within-task performances. Alternatively, the sample of mirror neurons studied might comprise a subset of mirror neurons characterized by perfect classification transfer across tasks, albeit restricted to particular time bins. In order to decide between these two scenarios, we compared the across-task discrimination performance of individual mirror neurons to the within-task performance. Rather than using the absolute discrimination level, we chose the difference relative to chance level for a particular time bin shown in Fig. 3A. As the concept of a *mirror mechanism* posits that the observation performance can be led back to an activation of a motor representation, we restricted this analytical step to a comparison of the exe → obs and the obs → obs discrimination performance.

Figure 4A depicts the resulting scatter plot of the exe → obs discrimination performance as a function of the obs → obs discrimination performance with individual data points representing individual neurons and time bins out of the 12 time bins covering the complete action sequence from start-button release until reward delivery. Data points indicating the same discrimination performance for within-task and across-task classifications would fall on the unity line (45 degrees line). On the other hand, a dissimilar discrimination performance would result in data points that are distributed around the exe → obs chance level (0 degrees line). If across-task performances were consistently weaker than within-task performances, we would expect the data points to be distributed around a regression line with a slope lower than 1. In fact, the regression line based on all data points had a slope of only 0.25 (Fig. 3A), indicating relatively weak transfer of decoding.

**Figure 4.**
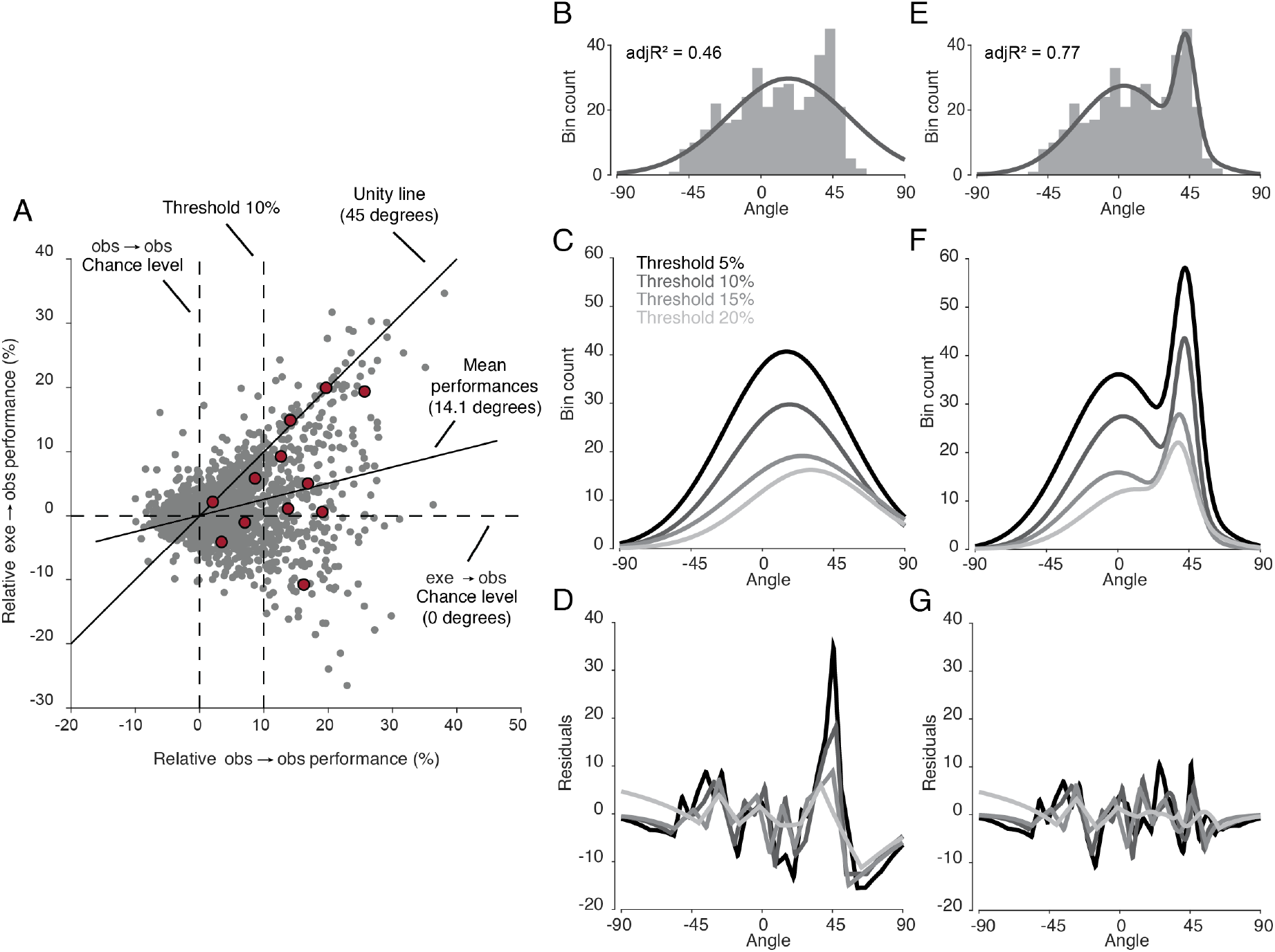
Identification of matching bins. A) Each dot in the scatter plot represents the exe → obs performance (related to the red curve in fig. 3A) and obs → obs performance (related to the green curve in fig. 3A) of a neuron in a time bin with respect to chance level. Each neuron contributed to this plot with 12 time bins between start-button release and reward. The red dots indicate the data points of an example neuron. Threshold 10% is one instance of the four filtering thresholds. The mean performances line (14.1 degrees) is a regression line of all the data points. B) The gray histogram shows the distribution of the angles of each dot above threshold 10% in A. The envelope depicts the single-Gaussian fit. C) Single-Gaussian fits for four different thresholds. D) The residuals of the single-Gaussian fits for the four thresholds. D) The two-Gaussian fit for threshold 10%. E) The two-Gaussian fits for the four thresholds. F) Two-Gaussian fits for the four thresholds. G) The residuals of the two-Gaussian fits for the four thresholds.

In order to assess how data points were distributed with respect to these reference lines, we transformed each data point to polar coordinates and created a distribution of angles. An obvious caveat is that meaningless data points were contained, characterized by the lack of discrimination performance in the reference case, i.e. the obs → obs classification. As there is no a priori criterion that would allow one to distinguish good discrimination performance from insufficient performance, we compared the influence of increasingly restrictive thresholds on the obs → obs performance (starting from 5% above chance level to 20% in steps of 5%).

As shown in Fig. 4B, the distribution of angles for a performance threshold of 10% clearly had a bimodal shape with a broad mode around zero and a narrower mode around 45°, the latter indicating a group of time bins with a complete match of classification performances. In accordance with the visual impression, trying to fit the distribution with a single Gaussian for all the thresholds (Fig. 4C) resulted in substantial residuals concentrated around 45 degrees, as shown in Fig. 4D, no matter which threshold was chosen (mean adjusted R^2^ across thresholds = 0.51). A two-Gaussian fit of the same distribution (10% threshold) resulted in a better fit that centered with a broad mode at 3.4 degrees and a second, narrower mode at 42.8 degrees (Fig. 4E). Figure 4F shows a consistent pattern of two-Gaussian fits (mean adjR^2^ = 0.76) across the four thresholds with robust means close to 0 and 45 degrees (inset), devoid of any concentration of residuals around particular angles (Fig. 4G). Hence, there is a distinct class of matching bins characterized by matched codes across the two tasks.

### A small subpopulation exists with a matched population code confined to the manipulation epoch

The question remains how the matching bins are distributed across neurons, and whether a subpopulation of neurons exists whose classification performances match throughout the action sequence. To this end, we separated matching bins from nonmatching bins, i.e. bins in which observed actions were discriminated by a matched code as opposed to a nonmatched code. To this end, we relied on the location of the trough between the two peaks of the two-Gaussian fit as the boundary between matching and nonmatching bins.

Considering the occurrence of matching and nonmatching bins in individual mirror neurons revealed four categories of neurons. Figure 5A distinguishes three of the four categories obtained for a performance threshold of 10%: a minority of “matching neurons” with only matching bins (10.5% of all mirror neurons), many more neurons with matching and non-matching bins (“mixed neurons”, 27.1%) and neurons with only nonmatching bins (29.4%). The fourth category (not shown in Fig. 5A) were neurons without any bin in which the observed actions could be dissociated (33.3%). Assuming that a biologically plausible readout would have to rely on the collective activity of distinct subsets of neurons, population classifications were carried out separately for each of these categories of neurons. For the subpopulation of neurons with only matching bins, we found a matched action code mainly during the manipulation epoch (Fig. 5B). The performance of exe → obs and obs → obs classifications started to be similar shortly before the actor touched the object and remained similar until the object was in its end position. After this point, however, the obs → obs performance remained high while the exe → obs performance gradually dropped to the chance level. In the subpopulation of mixed neurons (Fig. 5C) the exe → obs performance also increased during action observation, but it remained well below the obs → obs performance, indicating an only partially matching code. Not surprisingly, neurons with only nonmatching bins had an exe → obs population performance at chance level (Fig. 5D) and neurons that had neither a matching bin nor a nonmatching bin (i.e., did not discriminate the three actions during the observation task) showed a poor obs → obs performance (Fig. 5E). Hence, the population classification revealed that only a small subpopulation of mirror neurons exhibit a matched action code, which is mainly confined to the manipulation epoch.

**Figure 5.**
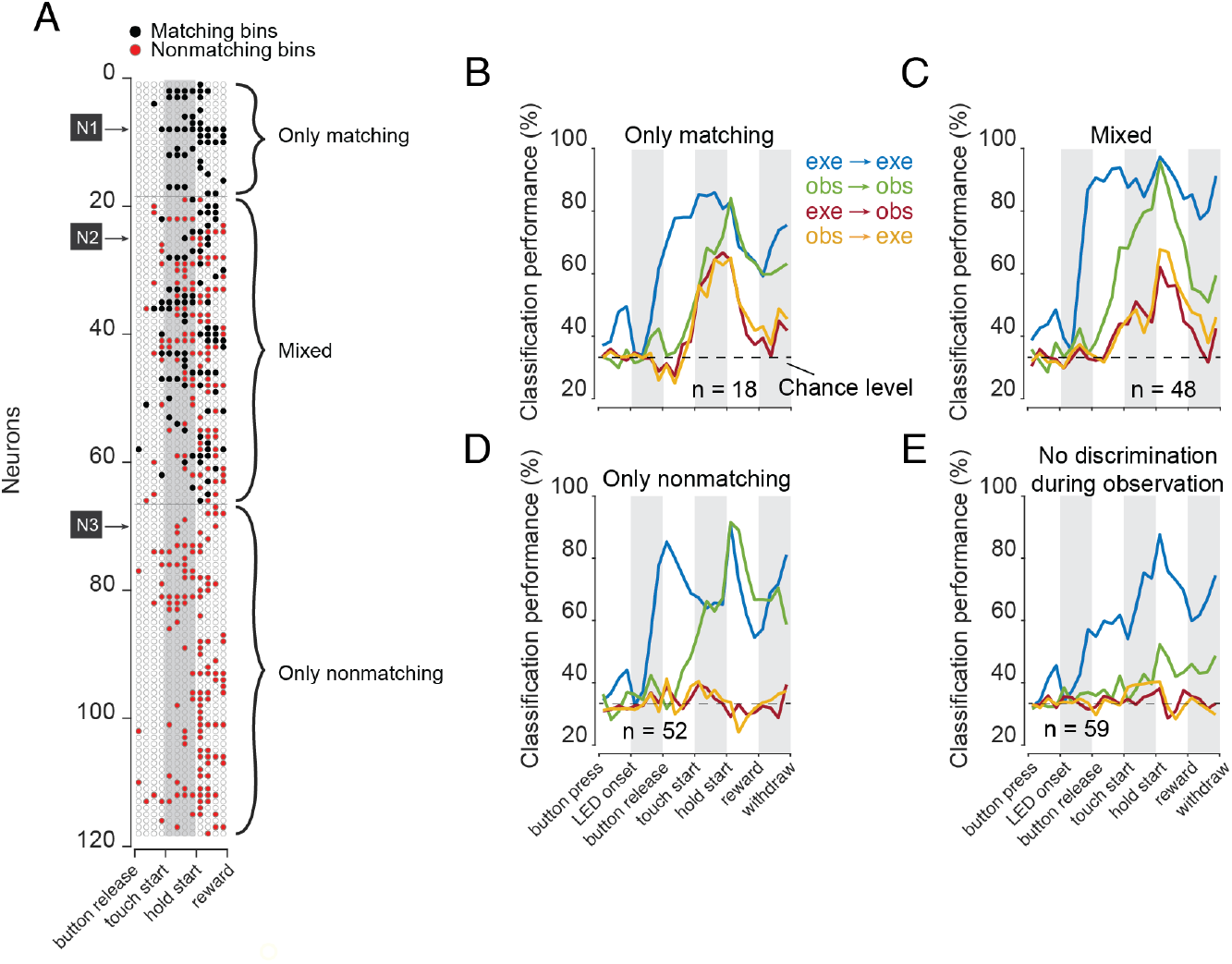
Subpopulations of mirror neurons. A) Mirror neurons that discriminated the three actions 10% better than chance (n = 118) are divided into three categories: neurons with only matching bins, mixed neurons with matching bins and nonmatching bins, neurons with only nonmatching bins. The arrows indicate the three exemplary neurons shown in figure 2. B) The population classification performance of the three subpopulations in A: B) neurons with only matching bins, C) neurons with matching and nonmatching bins, D) neurons with only nonmatching bin plus E) neurons that did not have any time bins with significant action discrimination during observation.

## Discussion

We report that three actions performed on visually identical objects could be decoded from the discharge rates of a population of F5 mirror neurons during both execution and observation. In both tasks, the decoding performance was optimal shortly after the end of the manipulation. A substantial degree of independence of the action codes for observation and execution is indicated by the fact that the ability of the action code for execution fell short of the quality of the one for observation when trying to predict the observed action. In other words, the level of action discrimination during observation achieved by the population of F5 mirror neurons, in its entirety, cannot be led back to a plain activation of a motor representation as assumed by the *mirror mechanism*. While not necessarily excluding a *mirror mechanism*, the ability to attain the necessary level of action discrimination during observation would in any case require supplementary analysis of visual inputs.

We considered the possibility that only a subpopulation of mirror neurons might fulfill the tenets of the mirroring concept. We could indeed unravel a small subset of mirror neurons that exhibited matched action codes for execution and observation for restricted epochs of the actions. We found neurons with matched action codes in one or more time bins in about one-third of the mirror neurons, with the number of bins per neuron varying. However, most of these neurons (73%; “mixed neurons”) also contained nonmatching bins, i.e., bins in which the observed actions were discriminated according to a code that differed from the one for execution. Correspondingly, when treated as a population acting collectively, these “mixed” mirror neurons were characterized by a code for observed actions that only partially matched the one for executed actions, again in contrast to the requirements of the *mirror mechanism*. Only a minority of 10.5% of mirror neurons contained matching bins for particular segments (i.e. well-defined relative time bins), but no discrimination of observed actions at other times. Population classification of this small subpopulation showed a matched action code mainly confined to the manipulation epoch. If this time restriction is disregarded, this subset of mirror neurons satisfies the requirements of the *mirror mechanism*, which leads the sensitivity to observed actions back to an activation of a motor representation of the same actions. Hence, by enabling a veridical activation of motor programs underpinning an agent’s own goal-directed actions, this subpopulation of mirror neurons might serve the understanding of others’ actions as suggested by Rizzolatti and Sinigaglia (Rizzolatti and Sinigaglia, 2016).

Yet, there are two caveats that complicate the acceptance of the conclusion. The first one is that any system for readout would need to know how to listen selectively to matching mirror neurons while ignoring all the others, unable to support veridical motor resonance. Certainly distinct anatomical or physiological features, if available, might allow the identification of matching mirror neurons. Our inability to identify such characteristic features may simply be a consequence of the well-known difficulty of extracellular recordings of neural activity from the brain of behaving animals to provide reliable information on the layer or type of a cortical neuron. The second caveat is that the readout mechanism would not only have to selectively listen to the subpopulation of matching neurons, but in addition be able to tune in to the right time epoch while ignoring others. However, how the epoch could be labeled in order to ensure the necessary temporal selection remains completely unclear. Hence trying to argue for a specific role of matching mirror neurons rather than taking them as outliers of a broad population of neurons sharing a common function does not seem to be particularly parsimonious.

We may note that the small proportion of 7.5% of all F5 neurons qualified as matching mirroring subpopulation in our study is in the same range as the proportion of “strictly congruent” mirror neurons (5.5%, 29/531) reported by Gallese and colleagues (Gallese, 1996) and the proportion of neurons with matched grip type preference reported recently by (Papadourakis, 2019). Although these numbers may not be directly comparable, considering differences between paradigms and analytical methods, the previous work concurs with our conclusion that only a small fraction of neurons may comply with the demands of a *mirror mechanism*. Moreover, the problem of how to filter out pertinent information from subpopulations lacking distinctive characteristics also pervades previous work trying to identify neurons compatible with the mirroring concept.

Trying to defend the existence of a small fraction of mirror neurons complying with the tenets of the mirroring concept inevitably leads to the question of what the function of all the other mirror neurons might be. The standard answer is that they are intermediate or hidden layer units in a network that tries to combine visual and motor information in order to accommodate the *mirror mechanism* (Elshaw et al., 2004; LeCun et al., 2015). This will certainly remain a valid possibility as long as we lack pertinent information allowing us to decide if matching mirror neurons have indeed distinct positions in the network or not. An alternative possibility is that the emergence of matching mirror neurons is an incidental and ultimately irrelevant consequence of a network that tries to associate visual information on observed actions with action commands. In order to allow visual information on observed actions to select and/or to shape action plans, matching would not be needed. An observed action is relevant for the observer and needs to be taken into account when weighing behavioral options. The notion that F5 mirror neurons might be instrumental in selecting and/or shaping actions and re-actions of course builds on the well-established role of the ventral premotor cortex in action control (Fogassi, 2001; Kraskov et al., 2009; Umilta et al., 2007; Wise et al., 1986).

In fact, a number of findings on action observation responses of F5 mirror neurons could be easily accommodated by the action selection framework. For instance, to select a response to an observed action, the angle from which the other is seen matters. Hence, it is not surprising that observation responses of mirror neurons have been shown to depend on the viewpoint (Caggiano, 2011). Action selection would benefit from anticipating actions chosen by the other. And indeed, the discharge of mirror neurons has been shown to reflect the expectation of an upcoming action (Maranesi et al., 2014), the expected final action goal the other is pursuing (Bonini et al., 2010), or the mere assumption of an ongoing action (Umiltà et al., 2001). The agent’s action choice will of course not only depend on information on the other’s action but to be useful it needs to depend on information on the relevance or value of the other’s action, including information on past experiences or information on the viability of potential action options. An example of the latter is the distance between the observer and the object, which has been demonstrated to modulate action observation responses of mirror neurons (Caggiano, 2009). As the distance of an object manipulated by the other is decisive for the question if it could be reached by the observer one may assume that a distance depending change of observation-related activity will also have an impact on the motor outflow from F5. During an action selection process that requires a decision between options, the respective value of competing options matters (Cisek, 2007). Assuming that the other’s actions influence the agent’s action choice, it is not surprising that action observation responses of mirror neurons are influenced by information on value (Caggiano et al., 2012; Pomper et al., 2020). One may correctly argue that all these observations are circumstantial, lacking the quality needed in order to firmly establish the action selection hypothesis based on irrefutabile causal evidence. However, if we believe that Ockham’s razor, i.e. to the principle of parsimony, has virtue in trying to solve a scientific problem, the action selection hypothesis may be taken as a useful pointer to a new, promising path towards a better understanding of a class of neurons that has remained enigmatic to the present day.

## Material and methods

### Experimental design

The experiments were performed on two adult male rhesus monkeys (Macaca mulatta). All experiments were approved by the local animal care committee (Regierungspräsidium Tübingen and Landratsamt Tübingen) and conducted in accordance with German and European law and the National Institutes of Health’s Guide for the Care and Use of Laboratory Animals, and carefully monitored by the veterinary service of the University of Tübingen.

### Behavioral tasks

Three objects to be manipulated by either the monkey or a human actor were fixed on a table and aligned parallel to the ground. The table rotated twice between trials (in the same or in opposite directions and for variable durations) to place randomly one of the three objects in front of the respective actor. The objects were metallic discs identical in shape and size (diameter 3.5 cm, height 0.5 cm). In the execution task, the center of the object at stake had a distance of 23 cm from the monkey’s eyes. In the observation task, in which a human standing opposite to the monkey acted on the relevant object, the distance to the monkey was 49 cm. Each object accommodated only one specific type of action: it could be either lifted (2.7 cm up), twisted (50° clockwise), or shifted (1.9 cm to the right). Note that the identical appearance of the three objects denied any clue on the type of action they accommodated.

In the execution task, the rotation of the table and the execution of the action on the target object by the monkey took place in darkness. In the observation task, a light was switched on when the object was rotated in place so that the monkey could observe the experimenter standing opposite to him (light was turned off with object release or if a trial was aborted).

A trial started when the object was rotated in place and the actor had pressed the start button. A variable time (1-1.5 s) after trial start, an LED next to the positioned object that had halted in front of the respective actor, was turned on for 0.1 to 0.3 s. The LED color indicated the manipulation to be performed: green or yellow for a lift, white or blue for a twist, red for a shift (in 5% of trials a yellow LED, noninformative for the type of object and located in a slightly different position was turned on; these trials were not analyzed here). The actor was allowed to release immediately the start-button to approach, manipulate, and finally hold the object against a resistance (against gravity for lift, against a spring for twist and shift). After a variable time after object hold start (0.3-0.8 s), the monkey — no matter if he had performed the action or had observed the human action — received water (or occasionally also units of a banana flavor high caloric drink) as reward, followed by the actor releasing the object. The amount of fluid per trial varied between 0.15 and 0.5 ml across sessions but was the same across the six actions (3 for each task) within a session.

The task events “start-button press”, “start-button release”, “LED onset”, “touch of object”, “start moving the object”, “reaching the hold position”, “reward”, and “withdraw from the object” by releasing it, were registered by mechanoelectrical sensors. A trial was aborted (and a beep was given as feedback to the monkey) if the start-button was released within 100 ms after LED onset (to motivate the monkey to use the LED), if the object was released before reward delivery or if a timeout was reached 10 s after trial start. To motivate the monkey even more to use the LED in the execution task, another timeout was active in 30% of trials (session-dependent rarely up to 100%) for the time period between “touch of object” to “start moving the object”: 0.15 s (rarely 0.1 s) for a twist and a shift, 0.35 s (rarely 0.3 s) for a lift). In the observation task, a trial was also aborted (and a beep was given as feedback to the monkey) if the monkey did not attend to the action as indicated by gaze not staying inside a given fixation window (see below). For the control of the experiment and the recording behavioral data, we deployed in-house open-source software (nrec: https://nrec.neurologie.uni-tuebingen.de, developed and maintained by F. Bunjes, J. Gukelberger, V. Gueler) running under Debian Linux on a standard PC.

### Measurement of eye movements

Eye position was recorded using an in-house video eye-tracker based on pupil detection in infrared light, operating at a sampling rate of 50 Hz, in one monkey and by permanently implanted search coils in the other monkey. Eye position recordings were calibrated at the beginning of each experimental session by asking the monkey to fixate a target dot displayed on a monitor, seen by the monkey in the table plane after redirection by a mirror in front of the monkey. In the table plane, the target dot (red color, 0.1° radius) appeared within the range the human actor performs his action, allowing the reliable association of object position and eye movement records. As the monkey’s head was painlessly fixed (see below) during the experiment, eye position corresponded to gaze position. In the action observation task, attention to the action was ensured by making the delivery of reward contingent on gaze staying within a fixation window of ±7° vertically and ±5° horizontally relative to the center of the relevant object.

### Surgical procedures

The animals were implanted with a titanium post accommodating the painless fixation of the head and a titanium recording chamber overlying area F5 of the left hemisphere. The correct position of the chamber was determined using information from a pre-surgical anatomical MRI scan. All surgical procedures were conducted under strict aseptic conditions deploying combination anesthesia with isoflurane (0.8%) and remifentanil (1–2 microgram/kg·min) with full control of relevant physiological parameters such as body temperature, heart rate, blood pressure, PO2, and PCO2 were monitored. Postoperatively, buprenorphine was given until signs of pain were gone. Animals were allowed to recover fully before starting the experiments.

### Electrophysiological recordings

Extracellular action potentials were recorded using glass-coated tungsten electrodes (0.5-to 2-MΩ impedance; Alpha Omega) using a multielectrode system equipped with up to eight probes (Alpha Omega Engineering). Action potentials of individual neurons were discriminated online resorting to template matching provided by Alpha Omega’s Multi Spike Detector.

The mirror neurons reported in this study were recorded from area F5 of the left hemispheres of the two experimental animals. Area F5 was targeted, guided by presurgical MRI, information on the location of the arcuate sulcus provided by electrode penetrations and a consideration of the response properties of neurons in F5 and in FEF as well as characteristic behavioral reactions to microstimulation with saccades evoked from the FEF and arm, hand, face, or mouth movements elicited when stimulating F5.

### Mirror neuron criteria

Except for the classification analysis to be described later, we used software based on MATLAB (2019a). For the analysis of discharge patterns, only well-isolated single units from area F5 were considered for which at least eight valid trials per condition were available. A trial was “valid” if the aforementioned sequence of task events (“start-button press” to “withdraw from the object”) was registered by the sensors without technical malfunction. A neuron was classified as an F5 mirror neuron resorting to established criteria (e.g., Pomper et al., 2020). To this end, a task was subdivided into 5 epochs: baseline (−750 ms to −250 ms from LED onset), approach phase (from start-button release until touching the object), manipulation phase I (from touching the object until moving the object), manipulation phase II (from moving the object until holding the object in its target position), hold phase (from holding the object in its target position until 150 ms later). A Friedman test with the factor “epoch” was performed for each action (lift, twist or shift) separately. A neuron had a motor or a visual response if the Friedman test was significant for at least one action (Bonferroni correction, alpha = .05/3) in execution or observation, respectively. If a neuron had a motor and a visual response, it was classified as a mirror neuron. Note that in accordance with standard procedures this classification did not require that the modulation affected the same epoch or same direction (i.e. discharge rate increase vs. decrease).

### Action preference analyses

Action preference was determined for each neuron and time bin separately for execution and observation based on three pairwise comparisons between two conditions (execution and observation) deploying Wilcoxon rank-sum tests: lift (L) vs. twist (T), lift (L) vs. shift (S), T vs. S, Bonferroni-corrected for the 3 comparisons and 24 time bins. An action was considered the preferred one if its discharge rate was significantly above the other two. If two actions were not statistically different but significantly above the third one, both were considered the preferred ones. In total, this resulted in three classes of narrow action preference (L, T, S) and another three classes of combined action preferences (LT, LS, TS). We finally asked if a mirror neuron had the same action preference for observation and execution.

### Classification analyses

We used a linear discriminant classifier (Fisher 1936) by running the ‘classify’ function available in MATLAB 2021a, in order to explore if the information offered by either single mirror neurons or the population discharge, predicted specific action types.

For a given time bin, the classification was performed on the average discharge rate of a neuron over the course of the time bin, using the same number of trials for each task condition. The number of trials for each task condition considered for single neuron classification was neuron-dependent and corresponded to the minimum number of trials per condition across both tasks and all conditions. For population classification, the number of trials for each task condition was eight, since only units with at least eight trials per condition were considered. The linear discriminant classifier was trained on 75% of trials selected randomly for one task (observation or execution) and the model obtained was then tested on the remaining trials of the same task, or in case of across-task classification, it was tested on all trials of the other task. For example, the model trained on discharge rate during execution was used in an across-task comparison to determine, how well it predicted the action from the discharge rate acquired during observation. The whole sequence of model calculation, within and across-task prediction, was repeated 200 times by randomly selecting new sets of training trials. The bin-wise means of the 200 iterations were taken as the performance of the classifier for individual neurons.

For the population classification, each neuron’s activity was used as a feature of the classifier. Six trials (75%) of a given task were used for training and the remaining two trials were used for testing. In the across-task classification, all eight trials of the other task were used for testing. The remaining procedure was like the single neuron classification.

## Acknowledgment

We acknowledge the contribution of Dan Arnstein in the design of the experiment and Dr. Peter Dicke for the presurgical MRT scans and implant design, assistance in surgeries and postsurgical care of animals as well as for his invaluable support in developing the experimental setup.

## Author contributions

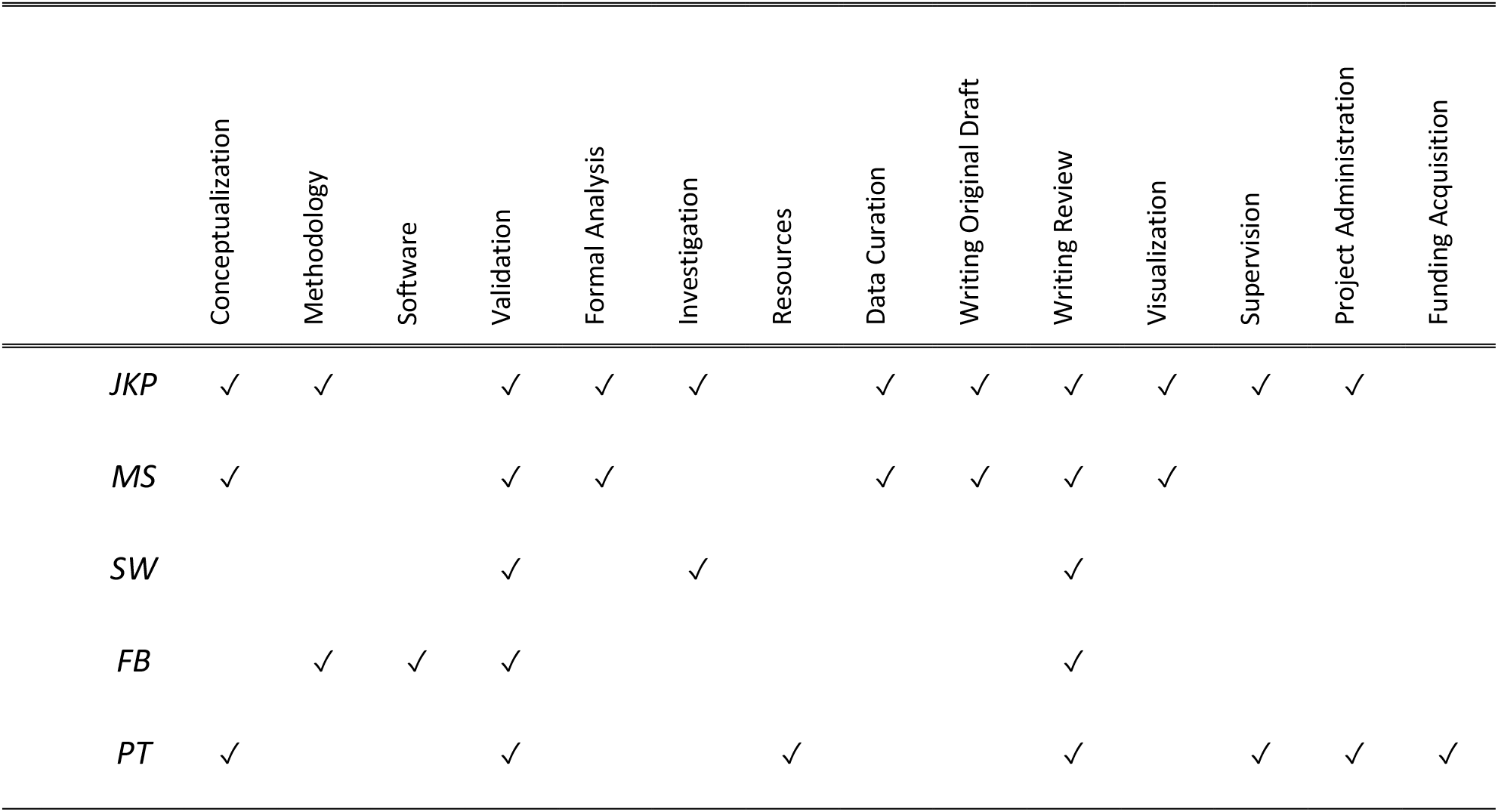

## References

Bonini L, Rozzi S, Serventi FU, Simone L, Ferrari PF, Fogassi L. 2010. Ventral premotor and inferior parietal cortices make distinct contribution to action organization and intention understanding. Cereb Cortex N Y N 1991 20:1372–1385. doi:10.1093/cercor/bhp200

Caggiano V, Fogassi L, Rizzolatti G, Casile A, Giese MA, Thier P. 2012. Mirror neurons encode the subjective value of an observed action. Proc Natl Acad Sci U S A 109:11848–11853. doi:10.1073/pnas.1205553109

Cook R, Bird G. 2013. Do mirror neurons really mirror and do they really code for action goals? Cortex J Devoted Study Nerv Syst Behav 49:2944–2945. doi:10.1016/j.cortex.2013.05.006

Csibra G. 2007. Action mirroring and action understanding: an alternative accountSensorimotor Foundations of Higher Cognition. Oxford: Oxford University Press. doi:10.1093/acprof:oso/9780199231447.003.0020

di Pellegrino G, Fadiga L, Fogassi L, Gallese V, Rizzolatti G. 1992. Understanding motor events: a neurophysiological study. Exp Brain Res 91:176–180. doi:10.1007/BF00230027

Elshaw M, Weber C, Zochios A, Wermter S. 2004. An associator network approach to robot learning by imitation through vision, motor control and language2004 IEEE International Joint Conference on Neural Networks (IEEE Cat. No.04CH37541). Presented at the 2004 IEEE International Joint Conference on Neural Networks. Budapest, Hungary: IEEE. pp. 591–596. doi:10.1109/IJCNN.2004.1379981

Ferrari PF, Rozzi S, Fogassi L. 2005. Mirror neurons responding to observation of actions made with tools in monkey ventral premotor cortex. J Cogn Neurosci 17:212–226. doi:10.1162/0898929053124910

Fisher RA. 1936.The Use of Multiple Measurements in Taxonomic Problems. 7:179–188.

Fogassi L. 2001. Cortical mechanism for the visual guidance of hand grasping movements in the monkey: A reversible inactivation study. Brain 124:571–586. doi:10.1093/brain/124.3.571

Fogassi L, Ferrari PF, Gesierich B, Rozzi S, Chersi F, Rizzolatti G. 2005. Parietal lobe: from action organization to intention understanding. Science 308:662–667. doi:10.1126/science.1106138

Gallese V, Fadiga L, Fogassi L, Rizzolatti G. 1996. Action recognition in the premotor cortex. Brain J Neurol 119 (Pt 2):593–609. doi:10.1093/brain/119.2.593

Hickok G. 2009. Eight problems for the mirror neuron theory of action understanding in monkeys and humans. J Cogn Neurosci 21:1229–1243. doi:10.1162/jocn.2009.21189

Kraskov A, Dancause N, Quallo MM, Shepherd S, Lemon RN. 2009. Corticospinal Neurons in Macaque Ventral Premotor Cortex with Mirror Properties: A Potential Mechanism for Action Suppression? Neuron 64:922–930. doi:10.1016/j.neuron.2009.12.010

LeCun Y, Bengio Y, Hinton G. 2015. Deep learning. Nature 521:436–444. doi:10.1038/nature14539

Maranesi M, Livi A, Fogassi L, Rizzolatti G, Bonini L. 2014. Mirror neuron activation prior to action observation in a predictable context. J Neurosci Off J Soc Neurosci 34:14827–14832. doi:10.1523/JNEUROSCI.2705-14.2014

Mukamel R, Ekstrom AD, Kaplan J, Iacoboni M, Fried I. 2010. Single-neuron responses in humans during execution and observation of actions. Curr Biol CB 20:750–756. doi:10.1016/j.cub.2010.02.045

Papadourakis V, Raos V. 2019. Neurons in the Macaque Dorsal Premotor Cortex Respond to Execution and Observation of Actions. Cereb Cortex 29:4223–4237. doi:10.1093/cercor/bhy304

Pomper JK, Spadacenta S, Bunjes F, Arnstein D, Giese MA, Thier P. 2020. Representation of the observer’s predicted outcome value in mirror and nonmirror neurons of macaque F5 ventral premotor cortex. J Neurophysiol 124:941–961. doi:10.1152/jn.00234.2020

Rizzolatti G, Camarda R, Fogassi L, Gentilucci M, Luppino G, Matelli M. 1988. Functional organization of inferior area 6 in the macaque monkey. II. Area F5 and the control of distal movements. Exp Brain Res 71:491–507. doi:10.1007/BF00248742

Rizzolatti G, Cattaneo L, Fabbri-Destro M, Rozzi S. 2014. Cortical mechanisms underlying the organization of goal-directed actions and mirror neuron-based action understanding. Physiol Rev 94:655–706. doi:10.1152/physrev.00009.2013

Rizzolatti G, Fadiga L, Gallese V, Fogassi L. 1996. Premotor cortex and the recognition of motor actions. Brain Res Cogn Brain Res 3:131–141. doi:10.1016/0926-6410(95)00038-0

Rizzolatti G, Sinigaglia C. 2016. The mirror mechanism: a basic principle of brain function. Nat Rev Neurosci 17:757–765. doi:10.1038/nrn.2016.135

Rizzolatti G, Sinigaglia C. 2010. The functional role of the parieto-frontal mirror circuit: interpretations and misinterpretations. Nat Rev Neurosci 11:264–274. doi:10.1038/nrn2805

Rochat MJ, Caruana F, Jezzini A, Escola L, Intskirveli I, Grammont F, Gallese V, Rizzolatti G, Umiltà MA. 2010. Responses of mirror neurons in area F5 to hand and tool grasping observation. Exp Brain Res 204:605–616. doi:10.1007/s00221-010-2329-9

Thompson EL, Bird G, Catmur C. 2019. Conceptualizing and testing action understanding. Neurosci Biobehav Rev 105:106–114. doi:10.1016/j.neubiorev.2019.08.002

Umilta MA, Brochier T, Spinks RL, Lemon RN. 2007. Simultaneous Recording of Macaque Premotor and Primary Motor Cortex Neuronal Populations Reveals Different Functional Contributions to Visuomotor Grasp. J Neurophysiol 98:488–501. doi:10.1152/jn.01094.2006

Umiltà MA, Kohler E, Gallese V, Fogassi L, Fadiga L, Keysers C, Rizzolatti G. 2001. I Know What You Are Doing: A Neurophysiological Study. Neuron 31:155–165. doi:10.1016/S0896-6273(01)00337-3

Wise SP, Weinrich M, Mauritz K-H. 1986. Movement-related activity in the premotor cortex of rhesus macaques. Progress in Brain Research. Elsevier. pp. 117–131. doi:10.1016/S0079-6123(08)63407-X

